# OCR-Stats: Robust estimation and statistical testing of mitochondrial respiration activities using Seahorse XF Analyzer

**DOI:** 10.1101/231522

**Authors:** Vicente A. Yépez, Laura S. Kremer, Arcangela Iuso, Mirjana Gušić, Robert Kopajtich, Eliška Koňaříková, Agnieszka Nadel, Leonhard Wachutka, Holger Prokisch, Julien Gagneur

**Affiliations:** Department of Informatics, Technical University Munich, Boltzmannstr. 3, 85748 Garching, Germany; Quantitative Biosciences Munich, Gene Center, Department of Biochemistry, Ludwig-Maximilians Universität München; Institute of Human Genetics, Helmholtz Zentrum München, Ingolstädter Landstr. 1, 85764 Neuherberg, Germany; Institute of Human Genetics, Klinikum rechts der Isar, Technical University Munich, Ismaninger Str. 22, 81675 München, Germany

**Keywords:** Oxygen Consumption Rate (OCR), mitochondrial respiration, bioenergetics, statistical testing, outlier detection

## Abstract

Accurate quantification of cellular and mitochondrial bioenergetic activity is of great interest in medicine and biology. Mitochondrial stress tests performed with Seahorse Bioscience XF Analyzers allow estimating different bioenergetic measures by monitoring oxygen consumption rates (OCR) of living cells in multi-well plates. However, studies of statistical best practices for determining OCR measurements and comparisons have been lacking so far. Therefore, we performed mitochondrial stress tests in 126 96-well plates involving 203 fibroblast cell lines to understand how OCR behaves across different biosamples, wells, and plates. We show that the noise of OCR is multiplicative, that outlier data points can concern individual measurements or all measurements of a well, and that the inter-plate variation is greater than intra-plate variation. Based on these insights, we developed a novel statistical method, OCR-Stats, that: i) robustly estimates OCR levels modeling multiplicative noise and automatically identifying outlier data points and outlier wells; and ii) performs statistical testing between samples, taking into account the different magnitudes of the between- and within-plates variations. This led to a significant reduction of the coefficient of variation across plates of basal respiration by 36% and of maximal respiration by 32%. Moreover, using positive and negative controls, we show that our statistical test outperforms existing methods, which either suffer from an excess of false positives (within-plates methods), or of false negatives (between-plates methods). Altogether, the aim of this study is to propose statistical good practices to support experimentalists in designing, analyzing, testing and reporting results of mitochondrial stress tests using this high throughput platform.

## 1. Introduction

Mitochondria are double membrane enclosed, ubiquitous, maternally inherited organelles present in most eukaryotic cells (1). They are mostly known as the powerhouses of the cell (2,3) due to their pivotal function in the cellular energy supply where ATP is generated by the mitochondrial respiratory chain in a process referred to as oxidative phosphorylation. Furthermore, mitochondria are involved in regulating reactive oxygen species (4), apoptosis (2), amino acid synthesis (5,6), cell proliferation (6), cell signaling (7), and in the regulation of innate and adaptive immunity (8). A decline in mitochondrial function, reflected by a diminished electron transport chain activity, is related to many human diseases ranging from rare genetic disorders (9) to common ones such as cancer (7,10), diabetes (11), neurodegeneration (12), and aging (3). One of the most informative tests of mitochondrial function is the quantification of cellular respiration, as it directly reflects electron transport chain impairment (9) and depends on many sequential reactions from glycolysis to oxidative phosphorylation (13). One of the last steps of cellular respiration is the oxidation of cytochrome c in complex IV which reduces oxygen to form water. Therefore, estimations of oxygen consumption rates (OCR) expressed in pmol/min, are conclusive for the ability to synthesize ATP and mitochondrial function, even more than measurements of intermediates (such as ATP or NADH) and potentials (16,17).

OCR was classically measured using a Clark-type electrode, which is time consuming, limited to whole cells in suspension and high yield, and does not allow automated injection of compounds (17). Also, experimentation with isolated mitochondria is ineffective because cellular regulation of mitochondrial function is removed during isolation (18). In the last few years, a new technology that calculates O_2_ concentrations from fluorescence (19) in a microplate assay format has been developed by the company Seahorse Bioscience (now part of Agilent Technologies) (20). It allows simultaneous real-time measurements of both OCR and ECAR in multiple cell lines and conditions, reducing the amount of required sample material and increasing the throughput (14,20).

Typically, OCR and ECAR are measured using the Seahorse XF Analyzer in 96 (or 24) well-plates at multiple time steps under three consecutive treatments (Fig. 1), in a procedure known as mitochondrial stress test (21). Under basal conditions, complexes I-IV exploit energy derived from electron transport to pump protons across the inner mitochondrial membrane. The thereby generated proton gradient is subsequently harnessed by complex V to generate ATP. Blockage of the proton translocation through complex V by oligomycin represses ATP production and prevents the electron transport throughout complexes I-IV due to the unexploited gradient, thus generating ATP-ase independent OCR only (Figs. 1A-B). Administration of FCCP, an ionophor, subsequently dissipates the gradient uncoupling electron transport from complex V activity and increasing oxygen consumption to a maximum level (Figs. 1A-B). Finally, mitochondrial respiration is completely halted using rotenone, a complex I inhibitor. There is still some remaining oxygen consumption that is independent from electron transport chain activity (Figs. 1A-B). This approach is label-free and non-destructive, so the cells can be retained and used for further assays (15).

**Figure 1.**
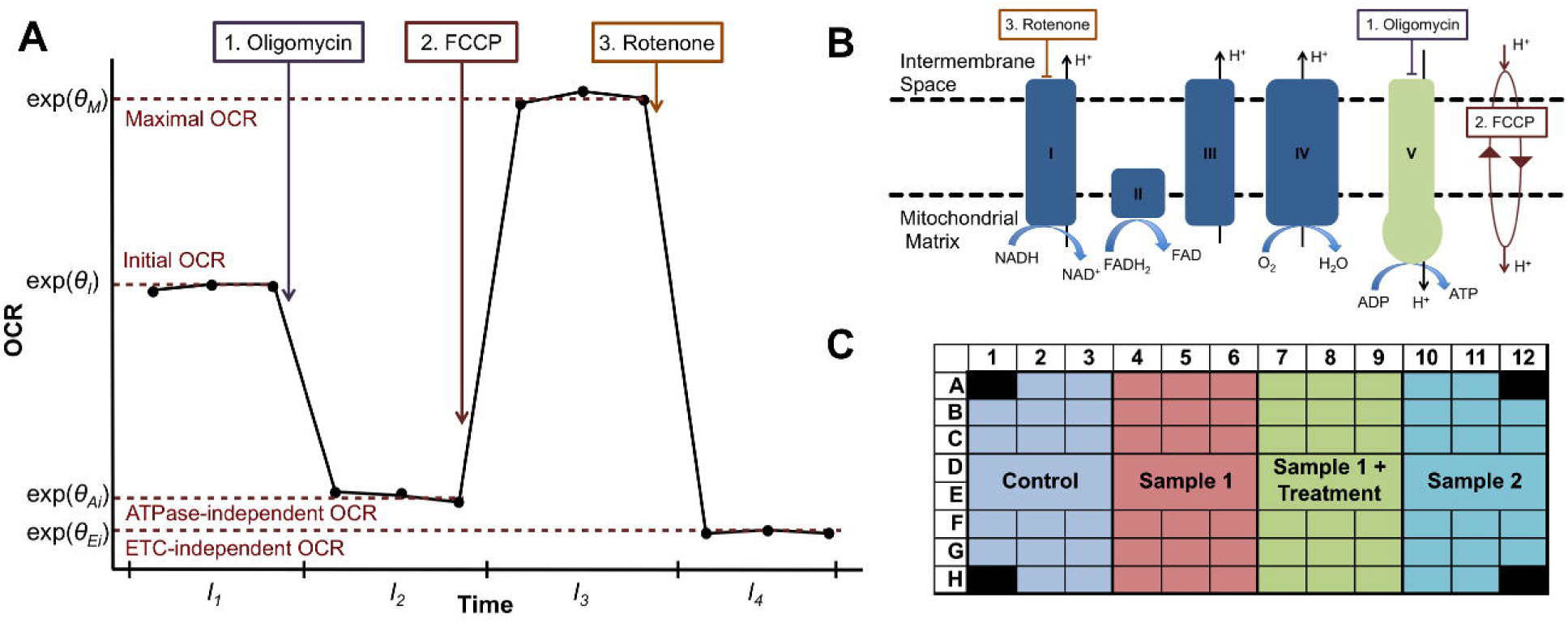
Principle of the mitochondrial stress test assay. (**A**) Cartoon illustration of OCR levels (y-axis) versus time (x-axis). Injection of the three compounds oligomycin, FCCP and rotenone delimit four time intervals within which OCR is roughly constant. (**B**) Targets of each compound in the electron transport chain. (**C**) Typical layout of a mitochondrial stress test 96-well plate.

OCR differences in the natural scale between the various stages of this procedure lead to the estimation of six different bioenergetics measures: basal respiration, proton leak, non-mitochondrial respiration, ATP-linked respiration, spare respiratory capacity, and maximal respiration (14,17) (Table 1). An increase in proton leak and a decrease in maximal respiration are indicators of mitochondrial dysfunction (17). ATP-linked respiration, basal respiration, and spare capacity alter also in response to ATP demand, which is not necessarily mitochondrion-related as it may be the consequence of deregulation of any cellular process altering general cellular energy demand.

**Table 1.** OCR ratios, abbreviations and equivalents

| OCR ratios | Abbreviation | Equivalent |
| --- | --- | --- |
| ETC-dependent OC proportion | E/I – proportion | Basal Respiration: $OCR_1 - OCR_4$ |
| ATPase-dependent OC proportion | A/I – proportion | ATP-linked respiration: $OCR_1 - OCR_2$ |
| ETC-dependent proportion of ATPase-independent OC | E/Ai - proportion | Proton Leak: $OCR_2 - OCR_4$ |
| Maximal OC fold change | M/I – fold change | Spare respiratory capacity: $OCR_3 - OCR_1$ |
| Maximal over ETC-independent OC fold change | M/Ei – fold change | Maximal respiration: $OCR_1 - OCR_4$ |
| Not defined as a ratio | Not defined | Non-mitochondrial respiration:<br>$OCR_4$ |

Current literature describing the Seahorse technology addresses experimental aspects regarding sample preparation (22,23), the amount of cells to seed (23,24), and compound concentration in different organisms (13,22,25). However, studies regarding statistical best practices for determining OCR levels and testing them against another are lacking. The sole definition of bioenergetic measures varies between authors, as well as the number of time points in each interval (usually three time points, but in some cases: one (26), two (27), and four or more (11)); and whether differences (6,13,28), ratios (12,29), or both (24,25) should be computed. Consequently, comparison of results across studies is difficult. Moreover, statistical power analyses for experimental design are often not provided. Differences in OCR between biosamples (e.g. patient vs. control, or gene knockout vs. WT) can be as low as 12 – 30% (30–32). Therefore, to design experiments with appropriate power to significantly detect such differences, it is important to know the source and amplitude of the variation within each sample, and reduce it as much as possible.

We performed and analyzed a large dataset of 126 experiments in 96-well plate format involving 203 different fibroblast cell lines, out of which 26 were seeded in more than one plate (Table S1). The large amount of between-plate and within-plate replicates allowed us to statistically characterize the nature and magnitude of biases and random variations in these data. We developed a statistical procedure called OCR-Stats, to extract robust and accurate oxygen consumption rates for each well, which translates into robust summarized values of the multiple replicates within and between plates. The OCR-Stats algorithm includes automatic outlier identification and controls for well and plates biases, which led to a significant increase in accuracy over state-of-the-art methods.

Between-well and between-plate biases, as well as random variations, were found to be multiplicative. This motivated us for a definition of bioenergetics measures based on ratios: ETC-dependent OC proportion, ATPase-dependent OC proportion, ETC-dependent proportion of ATPase-independent OC, and Maximal OC fold change (Table 1).

We provide estimators for each instance and show that they are empirically normally distributed. This permitted the use of linear regression models for assessing the statistical significance of bioenergetics measures comparisons between two biosamples. Using positive and negative controls from individuals known to have mitochondrial respiratory defects, we show that OCR-stats outperforms currently used statistical tests, which either suffer from an excess of false positives (within-plates methods), or of false negatives (between-plates methods).

Furthermore, our study provides experimental design guidance by i) showing that between-plate variation largely dominates within-plate variation, implying that it is important to seed the same biosamples in multiple plates, and ii) providing estimates of variances within and between plates for each bioenergetic measure allowing for statistical power computations. A free and pose source implementation of OCR-stats in the statistical language R is provided at github.com/gagneurlab/OCR-Stats.

## 2. Results

### 2.1 Experimental design and raw data

We measured OCR, ECAR, and cell number of 203 dermal fibroblast cultures derived from patients suffering from rare mitochondrial diseases and control cells from healthy donors (normal human dermal fibroblasts - NHDF, Methods, Table S1). These were assayed in 126 plates, all using the same protocol (Methods). Also, 26 cell lines were grown independently and measured in multiple plates. We will refer to these growth replicates as different biosamples. The NHDF cell line was seeded in all plates for assessment of potential systematic plate biases. The corners of each plate were left as blank, i.e. filled with media but not cells, to control for changes in temperature (22). One common layout of a plate is depicted in Fig. 1C, showing how each biosample is present in many well replicates. We seeded between 3 and 7 biosamples per plate (median = 4). This variation reflects typical set-ups of experiments in a lab performed over multiple years. Then, we used the standard mitochondrial stress test assay (21) leading to four time intervals with three time points each and denoted by Int_1_ (before adding any treatment), Int_2_ (after oligomycin), Int_3_ (after FCCP) and Int_4_ (after rotenone) (Fig. 1A). We also flagged wells that did not react as expected to the treatments and discarded them from the statistical analysis (Methods).

### 2.2 Random and systematic variations between replicates within plates

Representative replicate time series are shown in Fig. 2A, with data from 12 wells for one biosample in a single plate depicting commonly observed variations.

**Figure 2.**
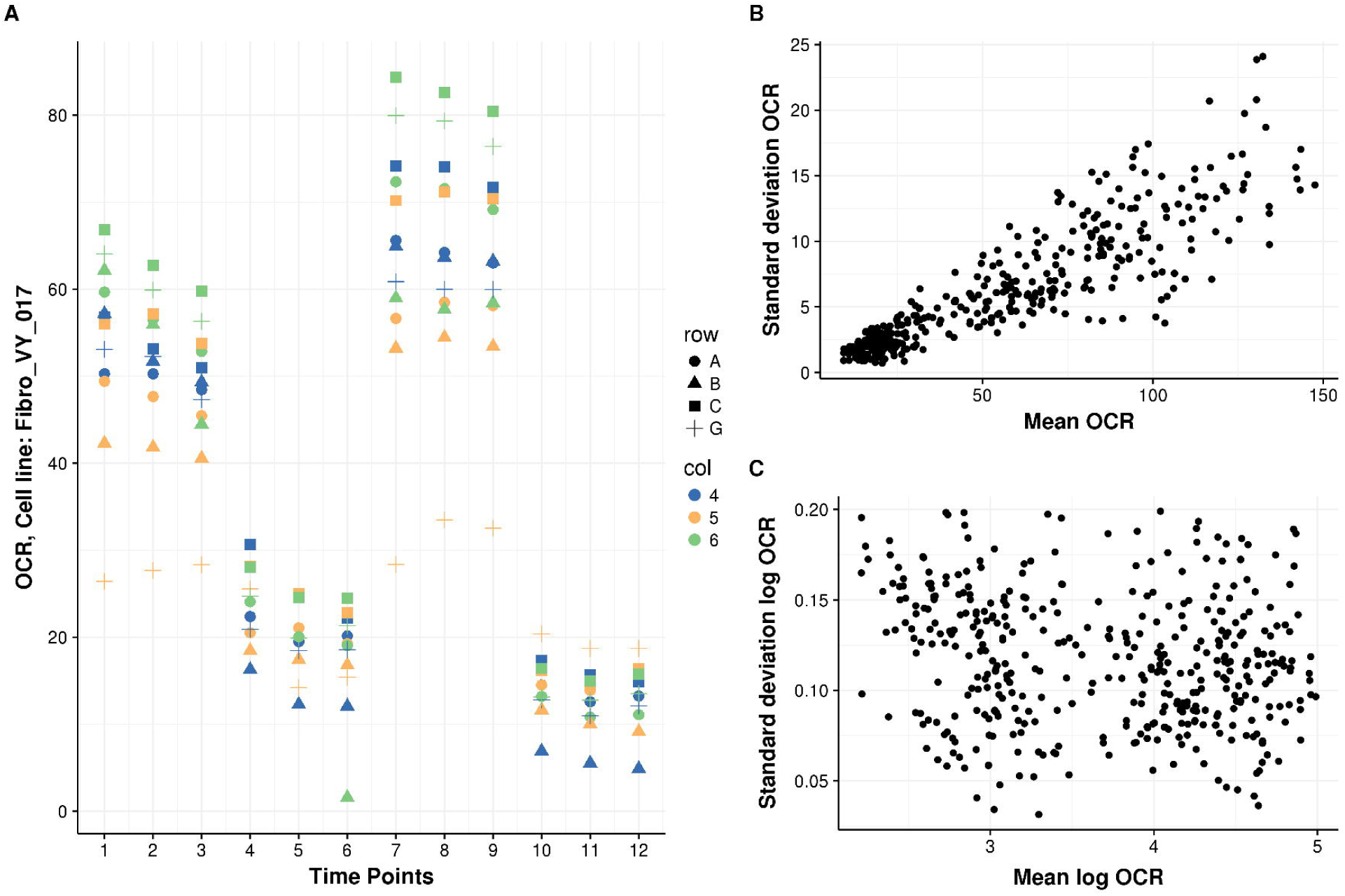
OCR behavior over time. (**A**) Typical time series replicates inside a plate. Behavior of OCR expressed in pmol/min (y-axis) of Fibro_VY_017 over time (x-axis). Colors indicate the row, and shape the column of 12 well replicates. Variation increases for larger OCR values, OCR has a systematic well effect and there exist two types of outliers: well-level and single-point. (**B**) Scatterplot of standard deviation (y-axis) vs. mean (x-axis) of all 3 time replicates of each interval, well and plate of OCR of NHDF only, shows a positive correlation (*n* = 409). (**C**) Same as (B) but for the logarithm of OCR, where the correlation disappears.

First, outlier data points occurred frequently. We distinguished two different types of outliers: entire series for a well (e.g., well G5 in Fig. 2A) and individual data points (e.g., well B6 at time point 6 in Fig. 2A). In the latter case, eliminating the entire series for well B6 would be too restrictive and result in losing valuable data from the other 11 valid time points. Therefore, methods to detecting outliers considering these two possibilities must be devised.

Second, we noticed a proportional dependence of OCR value and variance between replicates (Fig. 2B), suggesting that the error is multiplicative. Unequal variance, or heteroscedasticity, can strongly affect the validity of statistical tests and the robustness of estimations. We therefore propose modeling OCR on a logarithmic scale, where the dependency between variance and mean disappears (Figs. 2B, 2C). Respiratory chain enzyme activities such as NADH-ubiquinone reductase have also been shown to obey log-normal distributions (33).

Third, we observed systematic biases in OCR between wells (e.g., OCR values of well C6 are among the highest, while OCR values of well B5 are among the lowest at all time points, Fig. 2A). Variations in: cell number, initial conditions, treatment concentrations, or fluorophore sleeve calibration can lead to systematic differences between wells, which we refer to as well biases. To investigate whether well biases could be corrected using cell number to a large extend as in (26), we counted the number of cells after the experiments using Cyquant (Methods). As expected, median OCR for each interval grows linearly with cell number measured at the end of the experiment (Spearman rho between 0.32 and 0.47, *P* < 2.2e-16, Fig. S1A). However, the relation is not perfect reflecting important additional sources of variations and also possible noise in measuring cell number. Strikingly, dividing OCR by cell count led to a higher coefficient of variation (standard deviation divided by the mean) between replicate wells than without that correction (Fig. S1B). This analysis showed that normalization for cell number by division by raw cell counts is insufficient and motivated us to derive another method to capture well biases. Finally, we found that sex does not significantly associate with OCR levels (Fig. S2), in agreement with (34).

### 2.3 A statistical model of OCR

Building on these insights, we next introduced a statistical model for OCR within plates. For a given biosample in one plate, we modeled the logarithm of OCR y_w,t_ of well *w* at time point *t* as a sum of well biases, interval effects and noise, i.e.,:

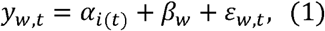

where α_*i(t)*_is the effect of the interval *i(t)* of time point *t*, *β_w_* is the relative bias of well *w* compared to a reference well, and ε_*w,t*_ is the error.

We defined the OCR levels *θ_i_* as the expected log OCR per interval, averaged over all wells:

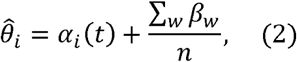

where *n* is the number of wells.

Note that the well bias is modeled independently for each plate, i.e., the bias of a certain well in one plate is different from the bias of the well at the same location in another plate.

We present now the OCR-Stats algorithm. For a given plate:

1. Fit the log linear model (1) using the least-squares method, which consists in minimizing ∑_*w*_∑_*t*_(y_*w,t*_-a_*i(t)*_- β_*w*_)_2_, thus obtaining the coefficients α_*i*_, β_*w*_. Compute 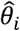 using (2).
2. For each time point *t* in interval *i* and well *w*, define the OCR residual: 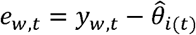, which is used to identify outliers (Methods, Fig. S3).
3. Identify and remove well level outliers, fit again, iteratively, until no more are found (Fig. S3A-B).
4. Identify and remove single point outliers, fit again, iteratively, until no more are found (Fig. S3C-D).
5. Scale back to natural scale in order to compute the bioenergetics measures (e.g.: Basal respiration = exp(θ_l_) - exp (θ_4_)), or take the ratio-based metrics (Tables 1 and 2).

**Table 2.** OCR ratio-based metrics

| OCR ratios | Metrics |
| --- | --- |
| ETC-dependent OC proportion | $\frac{OCR_1 - OCR_4}{OCR_1} = 1 - \exp(\theta_{Ei} - \theta_I)$ |
| ATPase-dependent OC proportion | $\frac{OCR_1 - OCR_2}{OCR_1} = 1 - \exp(\theta_{Ai} - \theta_I)$ |
| ETC-dependent proportion of ATPase-independent OC | $\frac{OCR_2 - OCR_4}{OCR_2} = 1 - \exp(\theta_{Ei} - \theta_{Ai})$ |
| Maximal OC fold change | $\frac{OCR_3}{OCR_1} = \exp(\theta_M - \theta_I)$ |
| Maximal over ETC-independent OC fold change | $\frac{OCR_3}{OCR_4} = \exp(\theta_M - \theta_{Ei})$ |

## 2.4 Variations within plates

We were then interested in determining the amplitude of the OCR variation between wells inside each plate, in order to compute the number of wells needed to obtain robust estimates 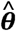. Using only the controls NHDF, we computed the standard deviation 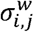 of the logarithm of OCR across all wells for each plate *j* and interval.Then, we computed the median across plates, thus obtaining one value 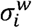 per interval 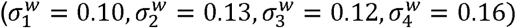. As we work in the logarithmic scale, the error in the natural scale becomes multiplicative and relative. The standard error of the estimates 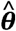 can be expressed as 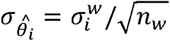, where *n*_*w*_ is the number of wells. The highest value of 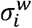 was 0.16, therefore cells should be seeded in 10 wells in order to get a relative error of 5%. This result is derived from variation after removing outliers, so considering that around 16.5% of wells were found to be outliers, ideally 10/(1 − 0.165) ≈ 12 wells should be used per biosample to get a relative error of 5%.

## 2.5 Variations between plates

After analyzing the OCR variation among wells inside plates, we set up to study the variation across multiple plates. Using data from the controls NHDF, we found that the variability between plates in all four intervals is much larger than between wells (Table S2, Fig. S4). We next asked whether a systematic plate bias exists that could be corrected for. We indeed observed a similar increase in OCR on interval 1 for both biosamples on plate #20140430 with respect to plate #20140428 (Fig. 3A). To test whether this tendency held across the repeated biosamples, we compared all replicate pairings with their respective NHDF controls and found a positive correlation (Fig. 3B). These differences can come from changes in temperature or the use of different sensor cartridges (13). Because the plate biases are systematic, they can be corrected for by using a log linear model (Methods). Nonetheless, the biases do not explain all the between plate variation, as the remaining variance is large (relative variance of the residuals: I_1_: 49.8%, I_2_: 51.6%, I_3_: 65.6% and I_4_: 55.9%). Therefore, when comparing two samples, it is important that they are seeded in the same plate, and that the test is performed multiple times.

**Figure 3.**
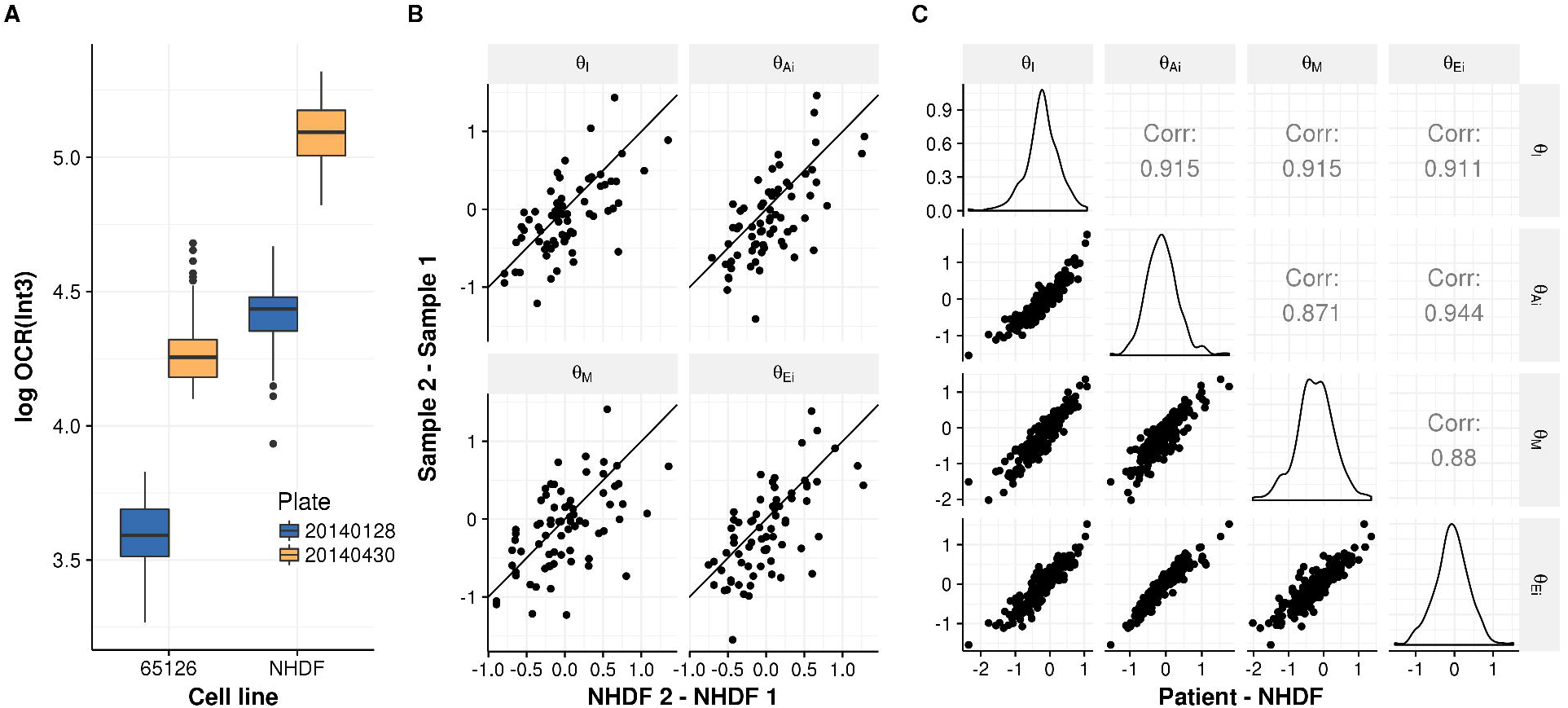
Plate bias. (**A**) Log of OCR in interval 3 (y-axis) for the cell lines #65126 and NHDF (x-axis) which were seeded in 2 different plates (color-coded). The similar increase in OCR from plate #20140128 to #20140430 in both biosamples suggests that there is a systematic plate bias. (**B**) Scatterplots of the differences of the logarithm of OCR levels θ of all possible 2 by 2 combinations of repeated biosamples across experiments (y-axis) against their respective controls (NHDF) (x-axis) show that there exists a positive correlation (I_1_: ρ= 0.64, *P* < 2.3×10^−8^, I_2_: ρ= 0.65, *P* < 1.2×10^−8^, I_3_: ρ= 0.52, *P* < 1.2×10^−5^, I_4_: ρ= 0.64, *P* < 1.4×10^−8^), confirming a systematic plate bias (*n* = 63). **(C)** Scatterplot of the difference of log OCR levels of patients vs. control NHDF (both axes) of every interval with respect to another. All intervals correlate with each other even after removing plate bias (by subtracting control values).

## 2.6 Statistical testing for the comparison of biosamples

In order to compare bioenergetics measures of two biosamples, we first need to evaluate the suitability of testing using differences versus testing using ratios of the OCR levels in the natural scale. As there is a remaining cell number effect after correcting for well biases (Fig. 3C), we recommend testing using ratios of OCR levels (or differences in the logarithmic scale) (Table 3).

**Table 3.**
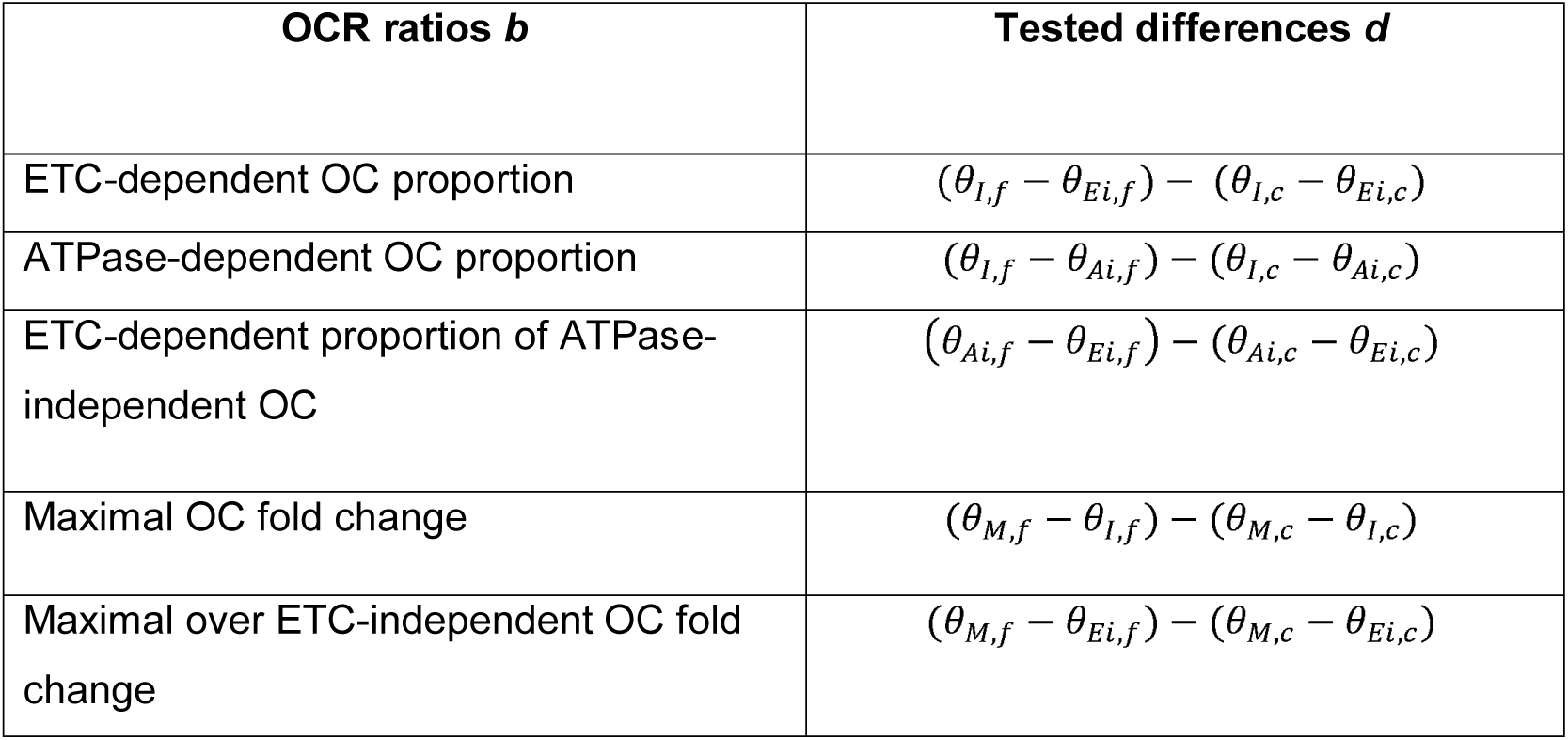
OCR ratio based differences for statistical testing

Subsequently, for any given OCR ratio *b* (eg. M/Ei - fold change), we test the differences of the log OCR ratios of a cell line *f* versus a control *c* using the following linear model:

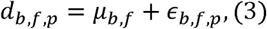

where *d*_*b*,*f*_,_*p*_corresponds to the difference of ratio *b* of a cell line *f* and the respective control on plate *p*. We solve it using linear regression, thus obtaining one value µ_*b,f*_per each ratio *b* and cell line *f*. We then compare these *µ*_*b,f*_values (which follow a t-Student distribution) against the null hypothesis µ_*b,f*_ =0 to compute p-values and confidence intervals (Figs. 4A, 4B, Methods).

**Figure 4.**
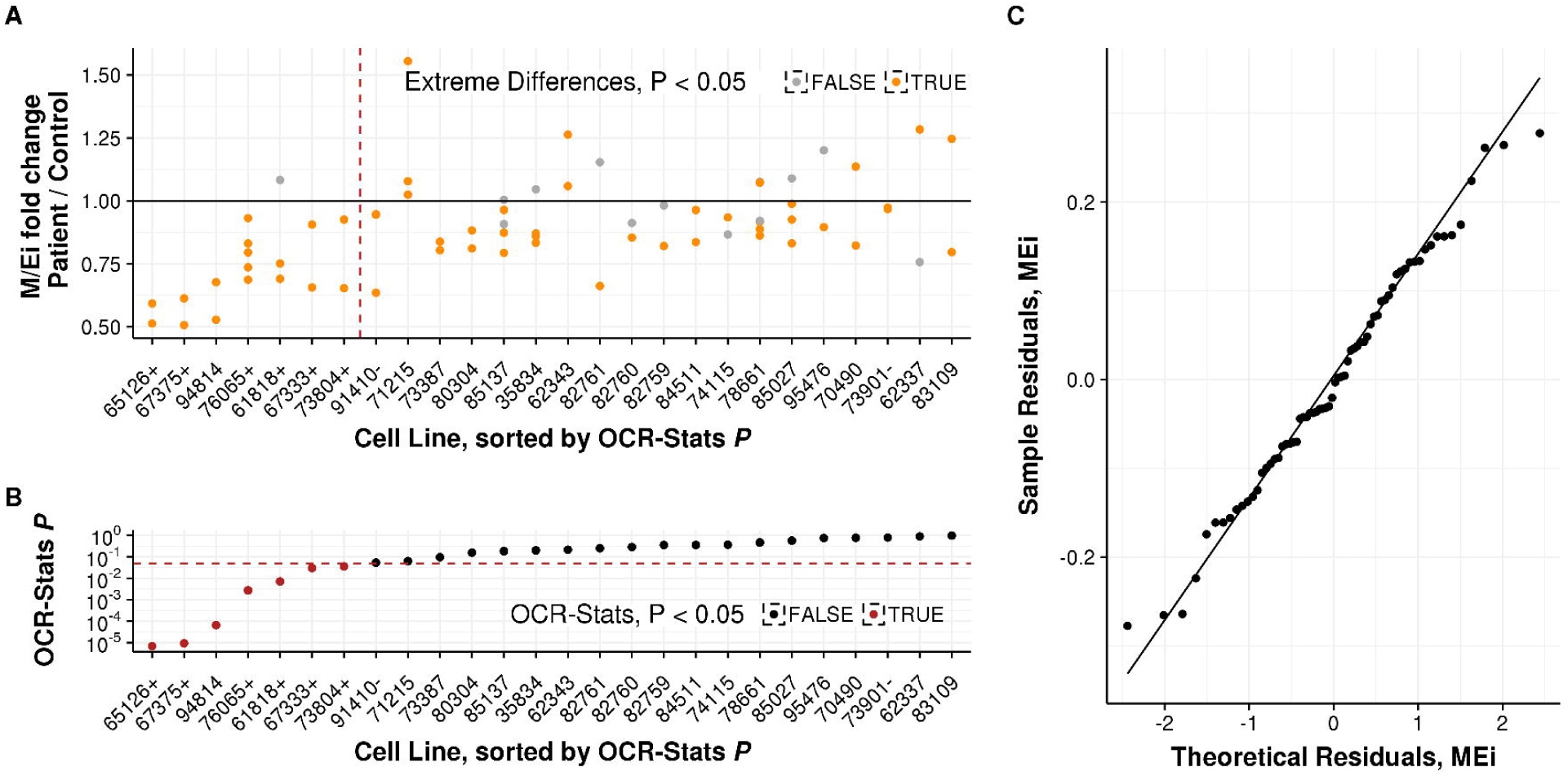
Statistical testing of M/Ei fold change patient vs. control on multiple plates. **(A)** Ratio of M/Ei fold change (y-axis) of all cell lines repeated across plates (x-axis) and their respective control, sorted by p-value obtained using the OCR-Stats method. Left of the red dashed line are cell lines with significantly lower M/Ei fold change using OCR-Stats. Dots in orange represent cell lines with significantly lower or higher M/Ei fold change using the ED method. Highlighted positive (+) and negative (-) controls. **(B)** Similar as (A), but depicting the p-value in logarithmic scale (y-axis) using OCR-Stats. Red dashed line at *P* = 0.05. Dots in red represent biosamples with significantly lower M/Ei fold change using the OCR-Stats method. **(C)** Quantile-quantile theoretical (x-axis) vs. observed (y-axis) plot of the residuals of the linear model (3) applied to M/Ei fold change.

## 2.7 Benchmark of OCR-Stats algorithm

In order to benchmark the OCR-Stats algorithm, we computed the coefficient of variation (standard deviation divided by mean) of the six bioenergetics measures in the natural scale of all repeated biosamples across plates for the following methods: i) the default Extreme Differences (ED) method (Methods) provided by the vendor, ii) the log linear (LL) corresponding to steps 1 and 2 of the OCR-Stats algorithm, iii) complete OCR-Stats (LL + outlier removal), and iv) OCR-Stats after correcting for plate effect (OCR-PE) using (4) (Methods).

Each step contributed to decreasing the coefficient of variation, obtaining a final significant reduction of 36% and 32% in basal and maximal respiration, respectively, from plate corrected OCR-Stats (OCR-PE) with respect to ED (*P* < 0.03, one-sided Wilcoxon test) (Fig. 5).

**Figure 5.**
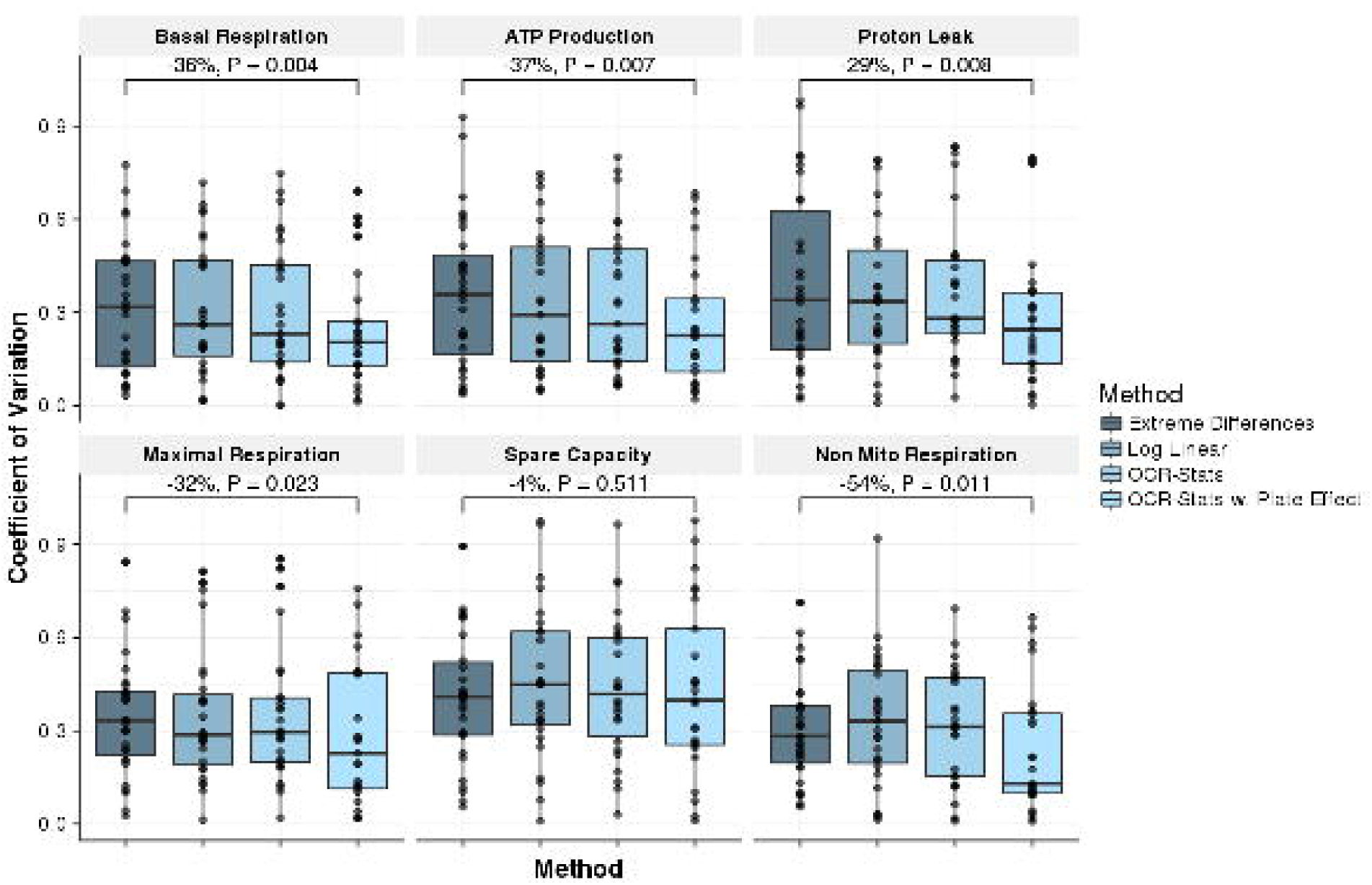
Benchmark using coefficient of variation. Coefficient of variation (CV = standard deviation / mean, y-axis) of replicates across experiments (*n*=26) using different methods (x-axis) to estimate the 6 bioenergetics measures. In all, except for Spare Capacity, OCR-Stats with plate effect showed significantly lower variation with respect to the Extreme Differences method. P-values obtained from one-sided paired Wilcoxon test.

## 2.8 Benchmark of OCR-Stats statistical testing method

We applied OCR-Stats statistical testing, Extreme Differences plus Wilcoxon test within each plate (within-plate ED), and Extreme Differences plus Wilcoxon test across plates (across-plate ED) to obtain the M/Ei ratio and maximal respiration (MR) of all the 26 cell lines that were seeded in more than one plate (Methods). For every approach, we computed p-values for significant fold changes against the controls. Six of these cell lines come from patients with rare variants in genes associated with an established cellular respiratory defect, allowing for assessing the sensitivity of each approach (Table S3, (35–39)). Additionally, two cell lines (#73901 and #91410) that showed no significant respiratory defects in earlier studies (40,41) served as negative controls.

The within-plate ED method reported significantly higher or lower MR for 56 out of 69 (81.2%) biosamples with respect to the control (Fig. 4A, Table S3). Moreover, all 26 cell lines had one or more significant biosamples on every plate, and 11 cell lines had one or more not significant sample (Fig. 4A). These ambiguous results show the importance of testing using multiple plates and advocate for a more robust approach than within-plate ED.

One approach to evaluate samples seeded in multiple plates is to perform a Wilcoxon test on the ED values averaged per plate (across-plate ED, Methods). However, this requires at least five plate replicates in order to obtain significant results. Here, only one cell line, #78661, was found significant this way. On this data, OCR-Stats was much more conservative than within-plate ED and found only 7 out of 26 (26.9%) cell lines to have aggregated significantly lower M/Ei than the control, including all 6 positive control cell lines (Figs. 4A, 4B, Table S3). Moreover, OCR-Stats did not report significant M/Ei differences for the two negative controls. There was no evidence against the normality and homoscedasticity assumption of OCR-Stats as the quantile-quantile plots of the residuals aligned well along the diagonal (Figs. 4C, S5). Altogether, these results show that OCR-Stats successfully identifies and removes variation within and between plates, providing more stable testing results which translates into less false positives.

## Discussion and conclusion

Mitochondrial studies using extracellular fluxes, specifically the XF Analyzer from Seahorse, are gaining popularity; therefore, it is of paramount importance to have a proper statistical method to estimate the OCR levels from the raw data. Here, we have developed such a model, the OCR-Stats algorithm, which includes approaches to control for well and plate biases, and automatic outlier identification. By doing so, we were able to significantly reduce the coefficient of variation of replicates across plates. Additionally, after analyzing the intra-plate variation, we suggest that the minimum number of wells replicates per biosample in a 96 well-plate should be 12.

We found that dividing cellular OCR by cell number was introducing more noise than was seen for uncorrected data. Here, we seeded always the same number of cells. Hence, the variations across wells that we observed in cell number at the end of the experiments are largely overestimated by noise in measurements. In other experimental settings in which different numbers of cells are seeded, we suggest to include an offset term to the model (1) equal to the logarithm of the seeded cell number to control for this variation by design. Also, the Seahorse XF Analyzer can be used on isolated mitochondria and on isolated enzymes, where a normalization approach is to divide OCR by mitochondrial proteins or enzyme concentration (42). However, as described here for cellular assays, robust normalization procedures require careful analysis.

We showed that there is roughly multiplicative bias between plates that can be controlled for to some extent by including control samples on every plate. To handle this plate bias, we proposed an extension of our within-plate robust linear regression approach adding a plate specific term. We demonstrated that OCR comparisons should be done using ratios rather than differences, as this eliminates sources of variation like cell number. We introduced a linear model, the OCR-Stats statistical testing, and showed that the results agree with previous results of patients diagnosed with mitochondrial disorders.

Significance with the OCR-Stats statistical algorithm can be reached by seeding a biosample in one plate only; provided there were other between-plate replicates to compute the inter-plate variance. Nevertheless, we still recommend performing at least 3 independent experiments of the same cell lines as one result alone can lead to wrong conclusions (Fig. 4A). Also note that a contaminated sample can increase the variability, affecting the significance of other samples. Therefore, it is important to detect them and discard them from further analysis.

In principle, OCR-Stats should be able to estimate ECAR levels. Similar analyses as performed here should be done beforehand in order to guarantee that the method is indeed applicable. Preliminary investigations suggest that the nature of noise (outliers, multiplicative) is similar than for OCR.

## Methods

### Biological material

All biosamples come from primary fibroblast cell lines of humans suffering from rare mitochondrial diseases, established in the framework of the German and European networks for mitochondrial disorders mitoNet and GENOMIT. The controls are primary patient fibroblast cell lines, normal human dermal fibroblasts (NHDF) from neonatal tissue, commercially available from Lonza, Basel, Switzerland.

### Measure of extracellular fluxes using Seahorse XF96

We seeded 20,000 fibroblasts cells in each well of a XF 96-well cell culture microplate in 80 ml of culture media, and incubated overnight at 37°C in 5% CO_2_. The four corners were left only with medium for background correction. Culture medium is replaced with 180 ml of bicarbonate-free DMEM and cells are incubated at 37°C for 30 min before measurement. Oxygen consumption rates (OCR) were measured using a XF96 Extracellular Flux Analyzer (21). OCR was determined at four levels: with no additions, and after adding: oligomycin (1 μM); carbonyl cyanide 4-(trifluoromethoxy) phenylhydrazone (FCCP, 0.4 μM); and rotenone (2 μM) (additives purchased from Sigma at highest quality). After each assay, manual inspection was performed on all wells using a conventionally light microscope. Wells for which the median OCR level did not follow the expected order, namely, median(OCR(Int_3_)) > median(OCR(Int_1_)) > median(OCR(Int_2_)) > median(OCR(Int_4_)), were discarded (977 wells, 10.47%). It is important to notice that other cell lines, or cell lines under certain conditions may not react as expected to the standard treatments, so they should not be discarded. We also excluded from the analysis contaminated wells and wells in which the cells got detached (461 wells, 4.94%, Methods). All the raw OCR data is available in Table S4.

### Cell number quantification

Cell number was quantified using the CyQuant Cell Proliferation Kit (Thermo Fisher Scientific, Waltham, MA, USA) according to the manufacturer’s protocol. In brief, cells were washed with 200 µL PBS per well and frozen in the microplate at −80°C to ensure subsequent cell lysis. Cells were thawed and resuspended vigorously in 200 µL 1x cell-lysis buffer supplemented with 1x CyQUANT GR dye per well. Resuspended cells were incubated in the dark for 5 min at RT whereupon fluorescence was measured (excitation: 480 nm, emission: 520 nm).

### Extreme Differences (default) Method to compute bioenergetics measures

On every plate independently, for each well, on interval 1 take the OCR corresponding to the last measurement, on intervals 2 and 4 take the minimum and on interval 3 the maximum OCR value (14). Then, do the corresponding differences to estimate the bioenergetics measures. Report the results per patient as the mean across wells plus standard deviation or standard error, separately for each plate.

### Outlier Removal

For each sample *s* and well *w*, compute the mean across time points of its squared residuals: 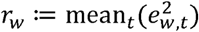 thus obtaining a distribution ***r***. Identify as outliers the wells whose *r*_*w*_> median(***r***) +5· mad(***r***), where mad, median absolute deviation, is a robust estimation of the standard deviation (Fig. S3A). We found that deviations by 5 mad from the median were selective enough in practice. Compute the vector of estimates 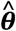 using the remaining wells and iterate this procedure until no more wells are identified as outliers. It required 8 iterations until convergence and around 16.5% of all the wells were found to be outliers (Fig. S3B).

Single point outliers are identified in a similar way. After discarding the wells that were found to be outliers in the previous step, categorize as outliers single data points whose 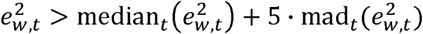 (Fig. S3C). Iterate until no more outliers are found. It required 19 iterations until convergence and approximately 6.1% of single points were found to be outliers (Fig. S3D).

### Plate effect model

In an attempt to correct for plate effect, we propose a log linear model where the levels er depend on interval *i*, samples *s* and plate *p*:

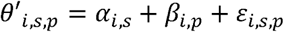

thus obtaining one coefficient β_*i,p*_for each plate-interval combination. These effects are added to the previous estimates: 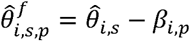 obtaining the final estimates 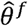 As for (1), the model is solved using linear regression.

For benchmarking, we cannot test using the estimates 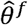, because we would fall into circularity, as correcting using β_*i,p*_forces replicates to have a closer value. Therefore, just for benchmarking purposes, we correct for plate effect using only the data from the controls NHDF *c* of each plate, namely:

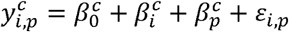

We solved it using linear regression and used the effects 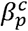 as offsets in (1). Then, we recomputed 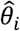 values accordingly. We scaled back to natural scale to calculate the bioenergetics measures and the coefficient of variation of all repeated biosamples (except the control).

### Multi-plate averaging method

In case of inter-plate comparisons, the multi-plate averaging methods takes the average and standard error of the bioenergetics measures obtained using the ED method of all repeated biosamples across plates (Agilent Technologies, 2016).

### OCR-Stats statistical testing

To evaluate the OCR ratios between a sample *f* and a control, both located on a plate *p*, we use the corresponding tested difference *d* (Table 3). We define 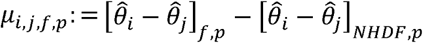 where *i* and *j* are any two different intervals. From there, we can obtain a t-statistic: 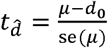, where *d*_*0*_ *= 0* as that is the value that we want to compare μ against, and *se* is the standard error. The t-statistic follows a t-distribution with *n* − *2* degrees of freedom, from which we can compute p-values. Moreover, we can obtain confidence intervals: 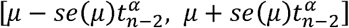 where (1 − α) is the confidence level and 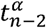 the (1 − α/2) quantile of the *t*_*n* − *2*_ distribution. Note that the normality assumption holds for the residuals E_*b,f,p*_(Figs. 4C, S5).

## Acknowledgements

We would like to thank Daniel Bader, Žiga Avsec, Jun Cheng and all members from the Gagneur Lab for valuable discussions and manuscript revision. This study was supported by the German Bundesministerium für Bildung und Forschung (BMBF) through the German Network for mitochondrial disorders (mitoNET, 01GM1113C to H.P.) E-Rare project GENOMIT (01GM1207 to H.P.), the Juniorverbund in der Systemmedizin ‘mitOmics’ (FKZ 01ZX1405A J.G., L.W. and V.A.Y.) and the DZHK (German Centre for Cardiovascular Research, L.S.K.). A Fellowship through the Graduate School of Quantitative Biosciences Munich (QBM) supports V.A.Y. H.P. is supported by EU FP7 Mitochondrial European Educational Training Project (317433). J.G., V.A.Y., L.S.K. and R.K. and H.P. are supported by EU Horizon2020 Collaborative Research Project SOUND (633974). We thank the Cell lines and DNA Bank of Pediatric Movement Disorders and Mitochondrial Diseases of the Telethon Genetic Biobank Network (GTB09003).

## Author contributions

J.G. and H.P. planned the project and overviewed the research. H.P. designed the experiments. V.A.Y. curated and analyzed the data. J.G. devised the statistical analysis. L.S.K., A.I., E.K., M.G., R.K., and A.N. performed the mitochondrial stress test experiments and cell number quantification. V.A.Y., L.W. and J.G. made the figures. V.A.Y. and J.G. wrote the manuscript. All authors performed critical revision of the manuscript.

## References

1. Gorman GS, Chinnery PF, DiMauro S, Hirano M, Koga Y, McFarland R, et al. Mitochondrial diseases. Nat Rev Dis Prim [Internet]. Macmillan Publishers Limited; 2016;2. Available from: http://www.nature.com/articles/nrdp201680

2. Bhola PD, Letai A. Mitochondria-Judges and Executioners of Cell Death Sentences. Mol Cell [Internet]. Elsevier Inc.; 2016;61(5):695–704. Available from: http://dx.doi.org/10.1016/j.molcel.2016.02.019

3. Sun N, Youle RJ, Finkel T. The Mitochondrial Basis of Aging. Mol Cell [Internet]. Elsevier Inc.; 2016;61(5):654–66. Available from: http://dx.doi.org/10.1016/j.molcel.2016.01.028

4. Wallace DC. Why do we still have a maternally inherited mitochondrial DNA? Insights from evolutionary medicine. Annu Rev Biochem [Internet]. 2007;76(1):781–821. Available from: http://www.annualreviews.org/doi/abs/10.1146/annurev.biochem.76.081205.150955

5. Birsoy K, Wang T, Chen WW, Freinkman E, Abu-Remaileh M, Sabatini DM. An Essential Role of the Mitochondrial Electron Transport Chain in Cell Proliferation Is to Enable Aspartate Synthesis. Cell [Internet]. 2015;162:540–51. Available from: http://linkinghub.elsevier.com/retrieve/pii/S0092867415008533

6. Sullivan LB, Gui DY, Hosios AM, Bush LN, Freinkman E, Vander Heiden MG. Supporting Aspartate Biosynthesis Is an Essential Function of Respiration in Proliferating Cells. Cell [Internet]. Elsevier Inc.; 2015;162(3):552–63. Available from: http://linkinghub.elsevier.com/retrieve/pii/S0092867415008545

7. Zong W-X, Rabinowitz JD, White E. Mitochondria and Cancer. Mol Cell [Internet]. Elsevier Inc.; 2016;166(3):555–66. Available from: http://dx.doi.org/10.1016/j.molcel.2016.02.011

8. Weinberg SE, Sena LA, Chandel NS. Mitochondria in the regulation of innate and adaptive immunity. Immunity [Internet]. Elsevier Inc.; 2015;42(3):406–17. Available from: http://dx.doi.org/10.1016/j.immuni.2015.02.002

9. Titov D V, Cracan V, Goodman RP, Peng J, Grabarek Z, Mootha VK. Complementation of mitochondrial electron transport chain by manipulation of the NAD+/NADH ratio. Science (80-) [Internet]. 2016;352(6282):231–5. Available from: http://www.ncbi.nlm.nih.gov/pubmed/27124460

10. Wallace DC. Mitochondria and cancer. Nat Rev Cancer [Internet]. Nature Publishing Group; 2012;12(10):685–98. Available from: http://www.nature.com/doifinder/10.1038/nrc3365

11. Dunham-Snary KJ, Sandel MW, Westbrook DG, Ballinger SW. Redox Biology A method for assessing mitochondrial bioenergetics in whole white adipose tissues. Redox Biol [Internet]. Elsevier Ltd.; 2014;2:656–60. Available from: http://dx.doi.org/10.1016/j.redox.2014.04.005

12. Yao J, Irwin RW, Zhao L, Nilsen J, Hamilton RT, Brinton RD. Mitochondrial bioenergetic deficit precedes Alzheimer’s pathology in female mouse model of Alzheimer’s disease. PNAS. 2009;106(34):14670–5.

13. Koopman M, Michels H, Dancy BM, Kamble R, Mouchiroud L, Auwerx J, et al. A screening-based platform for the assessment of cellular respiration in Caenorhabditis elegans. Nat Protoc [Internet]. Nature Publishing Group; 2016;11(10):1798–816. Available from: http://www.ncbi.nlm.nih.gov/pubmed/27583642

14. Divakaruni AS, Paradyse A, Ferrick DA, Murphy AN, Jastroch M. Analysis and interpretation of microplate-based oxygen consumption and pH data. Methods in Enzymology. 2014. 275-307 p.

15. Ferrick DA, Neilson A, Beeson C. Advances in measuring cellular bioenergetics using extracellular flux. Drug Discov Today. 2008;13(March).

16. Dmitriev RI, Papkovsky DB. Optical probes and techniques for O 2 measurement in live cells and tissue. Cell Mol Life Sci. 2012;69:2025–39.

17. Brand MD, Nicholls DG. Assessing mitochondrial dysfunction in cells. Biochem J. 2011;435(2):297–312.

18. Hill BG, Benavides GA, Jr JRL, Ballinger S, Italia LD. Integration of cellular bioenergetics with mitochondrial quality control and autophagy. Biol Chem. 2012;393(12):1485–512.

19. Gerencser AA, Neilson A, Choi SW, Edman U, Yadava N, Oh RJ, et al. Quantitative Microplate-Based Respirometry with Correction for Oxygen Diffusion. Anal Chem. 2009;81(16):6868–78.

20. Ribeiro SM, Giménez-cassina A, Danial NN. Measurement of Mitochondrial Oxygen Consumption Rates in Mouse Primary Neurons and Astrocytes. Methods Mol Biol. 2015;1241:59–69.

21. Agilent Technologies. Mito Stress Test Kit, User Guide. 2017;(103016–400). Available from: https://www.agilent.com/cs/library/usermanuals/public/XF_Cell_Mito_Stress_Test_Kit_User_Guide.pdf

22. Dranka BP, Benavides GA, Diers AR, Giordano S, Blake R, Reily C, et al. Assessing bioenergetic function in response to oxidative stress by metabolic profiling. Free Radic Biol Med. 2011;51(9):1621–35.

23. Zhang J, Nuebel E, Wisidagama DRR, Setoguchi K, Hong JS. Measuring energy metabolism in cultured cells, including human pluripotent stem cells and differentiated cells. Nat Protoc. 2012;7(6).

24. Zhou W, Choi M, Margineantu D, Margaretha L, Hesson J, Cavanaugh C, et al. HIF1α induced switch from bivalent to exclusively glycolytic metabolism during ESC-to-EpiSC/hESC transition. EMBO J [Internet]. Nature Publishing Group; 2012;31(9):2103–16. Available from: http://emboj.embopress.org/cgi/doi/10.1038/emboj.2012.71%5Cn http://www.pubmedcentral.nih.gov/articlerender.fcgi?artid=PMC3343469

25. Shah-Simpson S, Pereira CFA, Dumoulin PC, Caradonna KL, Burleigh BA. Bioenergetic profiling of Trypanosoma cruzi life stages using Seahorse extracellular flux technology. Mol Biochem Parasitol [Internet]. Elsevier B.V.; 2016;208(2):91–5. Available from: http://dx.doi.org/10.1016/j.molbiopara.2016.07.001

26. Dranka BP, Hill BG, Darley-Usmar VM. Mitochondrial reserve capacity in endothelial cells: The impact of nitric oxide and reactive oxygen species. Free Radic Biol Med [Internet]. Elsevier Inc.; 2010;48(7):905–14. Available from: http://dx.doi.org/10.1016/j.freeradbiomed.2010.01.015

27. Chacko BK, Kramer P a, Ravi S, Benavides G a, Mitchell T, Dranka BP, et al. The Bioenergetic Health Index: a new concept in mitochondrial translational research. Clin Sci [Internet]. 2014;127(6):367–73. Available from: http://www.ncbi.nlm.nih.gov/pubmed/24895057

28. Invernizzi F, D’Amato I, Jensen PB, Ravaglia S, Zeviani M, Tiranti V. Microscale oxygraphy reveals OXPHOS impairment in MRC mutant cells. Mitochondrion [Internet]. Elsevier B.V. and Mitochondria Research Society. All rights reserved; 2012;12(2):328–35. Available from: http://dx.doi.org/10.1016/j.mito.2012.01.001

29. Zhang J, Khvorostov I, Hong JS, Oktay Y, Vergnes L, Nuebel E, et al. UCP2 regulates energy metabolism and differentiation potential of human pluripotent stem cells. EMBO J [Internet]. Nature Publishing Group; 2011;30(24):4860–73. Available from: http://emboj.embopress.org/cgi/doi/10.1038/emboj.2011.401

30. Stroud DA, Surgenor EE, Formosa LE, Reljic B, Frazier AE, Dibley MG, et al. Accessory subunits are integral for assembly and function of human mitochondrial complex I. Nature [Internet]. Nature Publishing Group; 2016;538(7623):1–17. Available from: http://www.nature.com/doifinder/10.1038/nature19754%5Cn http://www.ncbi.nlm.nih.gov/pubmed/27626371

31. Mitsopoulos P, Chang Y-H, Wai T, König T, Dunn SD, Langer T, et al. Stomatin-like protein 2 is required for in vivo mitochondrial respiratory chain supercomplex formation and optimal cell function. Mol Cell Biol [Internet]. 2015;35(10):1838–47. Available from: http://www.pubmedcentral.nih.gov/articlerender.fcgi?artid=4405640&tool=pmcentrez&rendertype=abstract

32. Almontashiri NAM, Chen HH, Mailloux RJ, Tatsuta T, Teng ACT, Mahmoud AB, et al. SPG7 Variant Escapes Phosphorylation-Regulated Processing by AFG3L2, Elevates Mitochondrial ROS, and Is Associated with Multiple Clinical Phenotypes. Cell Rep [Internet]. The Authors; 2014;7(3):834–47. Available from: http://dx.doi.org/10.1016/j.celrep.2014.03.051

33. Hautakangas MR, Hinttala R, Rantala H, Nieminen P, Uusimaa J, Hassinen IE. Evaluating clinical mitochondrial respiratory chain enzymes from biopsy specimens presenting skewed probability distribution of activity data. Mitochondrion [Internet]. Elsevier B.V. and Mitochondria Research Society; 2016;29:53–8. Available from: http://dx.doi.org/10.1016/j.mito.2016.05.004

34. Kramer PA, Chacko BK, George DJ, Zhi D, Wei C-C, Dell’Italia LJ, et al. Decreased Bioenergetic Health Index in monocytes isolated from the pericardial fluid and blood of post-operative cardiac surgery patients. Biosci Rep [Internet]. 2015;35(4):e00237–e00237. Available from: http://bioscirep.org/cgi/doi/10.1042/BSR20150161

35. Hildick-Smith GJ, Cooney JD, Garone C, Kremer LS, Haack TB, Thon JN, et al. Macrocytic anemia and mitochondriopathy resulting from a defect in sideroflexin 4. Am J Hum Genet [Internet]. The American Society of Human Genetics; 2013;93(5):906–14. Available from: http://dx.doi.org/10.1016/j.ajhg.2013.09.011

36. Pronicka E, Piekutowska-Abramczuk D, Ciara E, Trubicka J, Rokicki D, Karkucińska-Więckowska A, et al. New perspective in diagnostics of mitochondrial disorders: two years’ experience with whole-exome sequencing at a national paediatric centre. J Transl Med [Internet]. BioMed Central; 2016;14(1):174. Available from: http://translational- medicine.biomedcentral.com/articles/10.1186/s12967-016-0930-9

37. Haack TB, Rolinski B, Haberberger B, Zimmermann F, Schum J, Strecker V, et al. Homozygous missense mutation in BOLA3 causes multiple mitochondrial dysfunctions syndrome in two siblings. J Inherit Metab Dis. 2013;36(1):55–62.

38. Kremer LS, Bader DM, Mertes C, Kopajtich R, Pichler G, Iuso A, et al. Genetic diagnosis of Mendelian disorders via RNA sequencing. Nat Commun [Internet]. 2017;8:15824. Available from: http://www.nature.com/doifinder/10.1038/ncomms15824

39. Van Haute L, Dietmann S, Kremer L, Hussain S, Pearce SF, Powell CA, et al. Deficient methylation and formylation of mt-tRNAMet wobble cytosine in a patient carrying mutations in NSUN3. Nat Commun [Internet]. Nature Publishing Group; 2016;7(May):12039. Available from: http://www.nature.com/doifinder/10.1038/ncomms12039

40. Powell CA, Kopajtich R, D’Souza AR, Rorbach J, Kremer LS, Husain RA, et al. TRMT5 Mutations Cause a Defect in Post-transcriptional Modification of Mitochondrial tRNA Associated with Multiple Respiratory-Chain Deficiencies. Am J Hum Genet [Internet]. The Authors; 2015;97(2):319–28. Available from: http://dx.doi.org/10.1016/j.ajhg.2015.06.011

41. Kremer LS, Distelmaier F, Alhaddad B, Hempel M, Iuso A, Küpper C, et al. Biallelic Truncating Mutations in TANGO2 Cause Infancy-Onset Recurrent Metabolic Crises with Encephalocardiomyopathy. Am J Hum Genet. 2016;98(2):358–62.

42. Seahorse Bioscience. Normalizing XF metabolic data to cellular or mitochondrial parameters, User Guide. 2014; Available from: http://hpst.cz/sites/default/files/attachments/appnote-normalizing-metabolic-data.pdf

43. Agilent Technologies. Multi-File XF Report Generator, User Guide. 2016; Available from: http://www.agilent.com/cs/library/usermanuals/public/Report Generator User Guide_Seahorse XF Cell Mito Stress Test_MultiFile_RevA.pdf

